# *Plasmodium* SAS4/CPAP is a flagellum basal body component during male gametogenesis, but is not essential for parasite transmission

**DOI:** 10.1101/2021.12.03.471138

**Authors:** Mohammad Zeeshan, Declan Brady, Robert Markus, Sue Vaughan, David Ferguson, Anthony A. Holder, Rita Tewari

## Abstract

The centriole/basal body (CBB) is an evolutionarily conserved organelle acting as a microtubule organising centre (MTOC) to nucleate cilia, flagella and the centrosome. SAS4/CPAP is a conserved component associated with BB biogenesis in many model flagellated cells. *Plasmodium*, a divergent unicellular eukaryote and causative agent of malaria, displays an atypical closed mitosis with an MTOC, reminiscent of the acentriolar MTOC, embedded in the nuclear membrane at most proliferative stages. Mitosis during male gamete formation is accompanied by flagellum formation: within 15 minutes, genome replication (from 1N to 8N) and three successive rounds of mitosis without nuclear division occur, with coordinated axoneme biogenesis in the cytoplasm resulting in eight flagellated gametes. There are two MTOCs in male gametocytes. An acentriolar MTOC located with the nuclear envelope and a centriolar MTOC (basal body) located within the cytoplasm that are required for flagellum assembly. To study the location and function of SAS4 during this rapid process, we examined the spatial profile of SAS4 in real time by live cell imaging and its function by gene deletion. We show its absence during asexual proliferation but its presence and coordinated association and assembly of SAS4 with another basal body component, kinesin8B, which is involved in axoneme biogenesis. In contrast its separation from the nuclear kinetochore marker NDC80 suggests that SAS4 is part of the basal body and outer centriolar MTOC residing in the cytoplasm. However, deletion of the SAS4 gene produced no phenotype, indicating that it is not essential for male gamete formation or parasite transmission through the mosquito.

## Introduction

Centriole/basal bodies (CBBs) are associated with the microtubule organising centre (MTOC) that nucleates cilia, flagella and centrosomes, and are conserved ancestral organelles in eukaryotes (Carvalho-Santos et al., 2011; Nabais et al., 2020). Centrioles and basal bodies (BBs) share structural features and BBs are mainly associated with flagella or cilia organisation, and extend to produce an axoneme (Marshall, 2008). The canonical view of CBB biogenesis is associated with their role in the cell cycle: centriole duplication occurs, and each is segregated to a daughter cell during mitosis. However, in some organisms BB biology is more diverse, for example where the centrioles or BBs form de novo or exhibit non-canonical biogenesis, as observed in Naegleria and in some parthenogenic insect eggs (Fritz-Laylin and Fulton, 2016; Nabais et al., 2020; Nabais et al., 2017). SAS4/SAS6 are ancestral core proteins involved in basal body biogenesis, as predicted by phylogenetic analysis (Carvalho-Santos et al., 2010; Hodges et al., 2010).

*Plasmodium spp*., the causative agents of malaria, are apicomplexan parasites transmitted by mosquito vectors. Asexual replication in *Plasmodium* is by atypical closed endomitosis, with remarkable plasticity in unconventional aspects of cell division during its complex life cycle. In a crucial stage for parasite transmission, sexually committed cells - the male and female gametocytes – are formed in the mammalian host and activated following ingestion by the female mosquito vector during its blood meal. Activation in the midgut results in gametogenesis and formation of extracellular female and flagellate male gametes over a period of 15 minutes (Sinden, 1991; Sinden et al., 2010). During male gametogenesis within 8 minutes there are three rounds of genome replication from haploid (1N) to octaploid (8N) without nuclear division and this is followed by karyokinesis and cytokinesis resulting in eight haploid flagellated gametes in a process known as exflagellation (Sinden et al., 2010). Flagella assembly is very rapid and atypical, occurring within 15 minutes and without intra-flagellar transport (IFT) (Sinden, 1991; Sinden et al., 2010). There are no flagella at other stages of the life cycle and hence no clear centriole or basal body is observed at other proliferative life cycle stages. Clear centrioles with 9+1 or 9+2 microtubules can only be seen in flagella biogenesis during male gamete formation in *Plasmodium* (Sinden et al., 2010; Straschil et al., 2010; Zeeshan et al., 2019a). So-called centriolar plaques located within the nuclear envelope were described during nuclear division in asexual stages of proliferation and appears to serve as an MTOC in nuclear spindle formation (Arnot et al., 2011; Gerald et al., 2011). Centrin has been mapped to these plaques and used as a marker for the MTOC to follow the asynchronous replication dynamics during asexual replication. However, no centriole is present and these plaques resemble an acentriolar MTOC organising hemispindle (Bertiaux et al., 2021; Mahajan et al., 2008; Roques et al., 2019; Simon et al., 2021). In male gametocytes there are two MTOCs: an inner acentriolar MTOC located with the nuclear envelope and outer centriolar basal bodies located within the cytoplasm that are required for flagellum assembly. The marker, NDC80 shows that kinetochores are clustered in the nucleus in asexual stages with a rod like structure at spindle formation during the three successive rounds of genome replication during in gametogenesis that can be differentiated from the outer MTOC (Zeeshan et al., 2020b).

BB structure and the real time dynamics of its formation during this accelerated genome replication and chromosome segregation remain unclear in Plasmodium, although earlier ultrastructural studies by electron microscopy (EM) suggested atypical BB structure and flagella. (Sinden et al., 1976). Ultrastructure studies also confirmed that the nuclear envelope remains intact during the closed mitosis, so events in the nucleus and cytoplasm have to be coordinated for flagellated gamete formation to occur: the two MTOCs need to be organised and coordinated for both these compartments so that each microgamete receives a single flagellum. In recent studies we have shown that some molecular motors like Kinesin-8X and Kinesin-5 are associated with the spindle in the nuclear compartment (Zeeshan et al., 2020a; Zeeshan et al., 2019b) and others like Kinesin-8B and Kinesin-X4 are involved in axoneme biogenesis (Zeeshan et al., 2019a). Therefore the MTOC during male gametogenesis may be described as composed of an outer centriolar BB, which organises the axoneme and an inner acentriolar MTOC similar to the spindle pole body of yeast from where the mitotic spindle and chromosome segregation is organised (Zeeshan et al., 2019a; Zeeshan et al., 2019b). An earlier study with the basal body marker SAS6 showed it was present outside the nucleus in the cytoplasmic BB compartment during male gametogenesis and its deletion ablated male gametogenesis, blocking parasite transmission (Marques et al., 2015). Recently we and others have shown the dynamic profile of Kinesin-8B, another BB marker, in axoneme assembly and its deletion leads to disruption of axoneme assembly and loss of flagellum formation (Depoix et al., 2020; Zeeshan et al., 2019a).

Since SAS4 and SAS6 are both ancestral components of BB formation (Carvalho-Santos et al., 2011; Carvalho-Santos et al., 2010; Hodges et al., 2010), here we have investigated the real time temporal profile of SAS4 during male gametogenesis, particularly during the 8 minutes of rapid genome replication following activation. We also investigated its subcellular association with kinetochore and axoneme biogenesis by genetically crossing SAS4-green fluorescent protein (GFP) and NDC80-mCherry or Kinesin-8B-mCherry transgenic parasite lines to obtain male gametocytes expressing both fluorescent markers. To examine the role of SAS4 we generated SAS4 gene knockout (KO) parasites. The results reveal a de novo rapid synthesis of SAS4 and suggest an association with the outer centriolar BB as a doublet during male gametogenesis. However, in contrast to the SAS6 KO mutant, the SAS4 KO mutant developed normally indicating that this protein is not essential for parasite growth in the mosquito or transmission.

## Results

### Live cell imaging of SAS4-GFP reveals de novo basal body formation and its rapid dynamics during male gametogenesis

In order to study the expression and location of SAS4 during the *Plasmodium* life cycle, we generated a SAS4-GFP transgenic *P. berghei* line expressing the protein with a C-terminal GFP tag, by inserting an in-frame *gfp* coding sequence at the 3’ end of the endogenous *sas4* locus using single homologous recombination (Fig S1A). PCR analysis of genomic DNA using locus-specific diagnostic primers indicated correct integration of the GFP tagging construct (Fig S1B). This transgenic line was used to examine the spatiotemporal profile of SAS4-GFP protein expression and location by live cell imaging. By microscopy, SAS4 was not detectable in asexual blood stages but was located in the cytoplasm of male gametocytes (Fig 1). Therefore, we investigated its expression and location throughout male gametogenesis. Male gametogenesis is a rapid process of three rounds of genome replication, de-novo basal body formation and axoneme assembly followed by emergence of eight flagellated male gametes, known as exflagellation, that completes within 12-15 minutes (Sinden et al., 2010, Zeeshan, 2019 #114).

**Fig 1.**
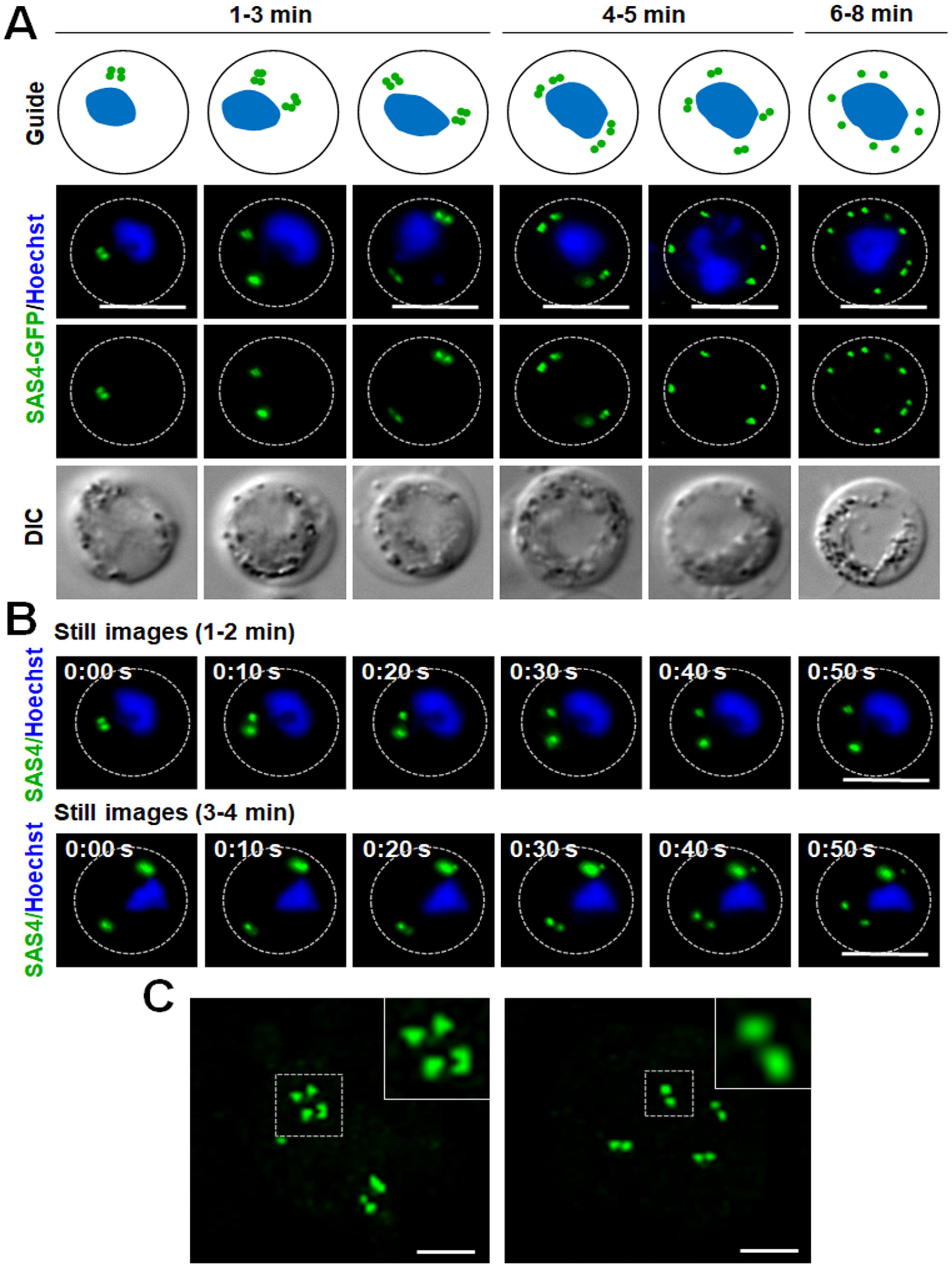
Localization of SAS4-GFP during male gametogenesis. **(A)** Live cell imaging of SAS4-GFP expression and location during endoreduplicative mitotic division in male gametogenesis. Scale bar = 5 μm. Schematic guide showing SAS4-GFP location in relation to nucleus (DNA) during male gametogenesis **(B)** Time-lapse screenshots for SAS4-GFP localization in male gametocytes at 1-2 min and 3-4 min post activation during male gametogenesis. Scale bar = 5 μm. **(C)** Super-resolution 3D imaging for SAS4-GFP localization in gametocytes fixed at 2- and 4-min post activation. Scale bar = 1 μm.

Live cell images showed multiple discrete SAS4-GFP foci in the cytoplasm of developing gametocytes, with the number of foci depending on the length of time after activation (Fig 1A). Within one minute post activation (mpa) of the gametocyte, four closely associated foci forming an SAS4-GFP tetrad were observed in the cytoplasm at one side of the nucleus (Fig 1A). The SAS4-GFP tetrad split later into two halves that moved apart to opposite sides of the nucleus within 3 mpa, still retaining the cytoplasmic location (Fig 1A, B and video S1). The two SAS4-GFP tetrads each split again into two doublets of SAS4-GFP and separated from each other within 4-5 mpa (Fig 1A, C and video S2). A final round of splitting and separation occurred to produce eight discrete SAS4-GFP foci (Fig 1A). A schematic diagram of this process is provided in the upper panel of Figure 1A.

To resolve further the SAS4-GFP foci after male gametocyte activation, 3D-structured illumination microscopy (SIM) was performed on fixed gametocytes expressing SAS4-GFP. The SIM images clearly showed two tetrads of SAS4-GFP in gametocytes at 2 mpa and four doublets by 4 mpa (Fig 1 D) indicating that SAS4 is present in closely apposed doublets.

### SAS4 associates with Kinesin-8B, a molecular motor that regulates basal body formation and axoneme assembly

Recently we showed that Kinesin-8B associates with the BBs and axonemes during *Plasmodium* male gametogenesis (Zeeshan et al., 2019a); live cell imaging showed the association of Kinesin-8B with the tetrad of BBs that serve as a template for axoneme assembly (Zeeshan et al., 2019a). To establish whether SAS4 is part of the BB and associated with tetrad of BB formation and axoneme assembly, we examined its location compared with that of Kinesin-8B. A parasite line expressing both SAS4-GFP and Kinesin-8B-mCherry was produced and used for live cell imaging by fluorescence microscopy to establish the spatiotemporal relationship of these two proteins. Within 1 mpa, the SAS4-GFP tetrad was observed at the centre of four Kinesin-8B-mCherry foci in the cytoplasm at one side of the nucleus (Fig 2A). Within 3 mpa we observed the duplication and separation of tetrads of SAS4 and Kinesin-8B (Fig 2A). To further resolve this dissociation, we performed 3D-SIM on fixed gametocytes expressing these two proteins. 3D-SIM images clearly show the SAS4 tetrads at the centre of kinesin-8B tetrads (Fig 2A, right hand panel). After arrival at either side of the nucleus, the emergence of axonemes was observed, as revealed by Kinesin-8B that later is only associated with axonemes (Fig 2A). The SAS4-GFP tetrad later split into doublets, which remain associated with growing axonemes labelled with Kinesin-8B-mCherry during their further split and separation (Fig 2A). At the end of the process, we observed eight SAS4-GFP foci associated with fully assembled axonemes (Fig 2A). These data shows that SAS4 is associated with kinesin-8B during a very early stage of basal body formation and remains associated with it throughout axoneme assembly during the rest of male gametogenesis. A schematic diagram of this process is provided in Fig 2B.

**Figure 2.**
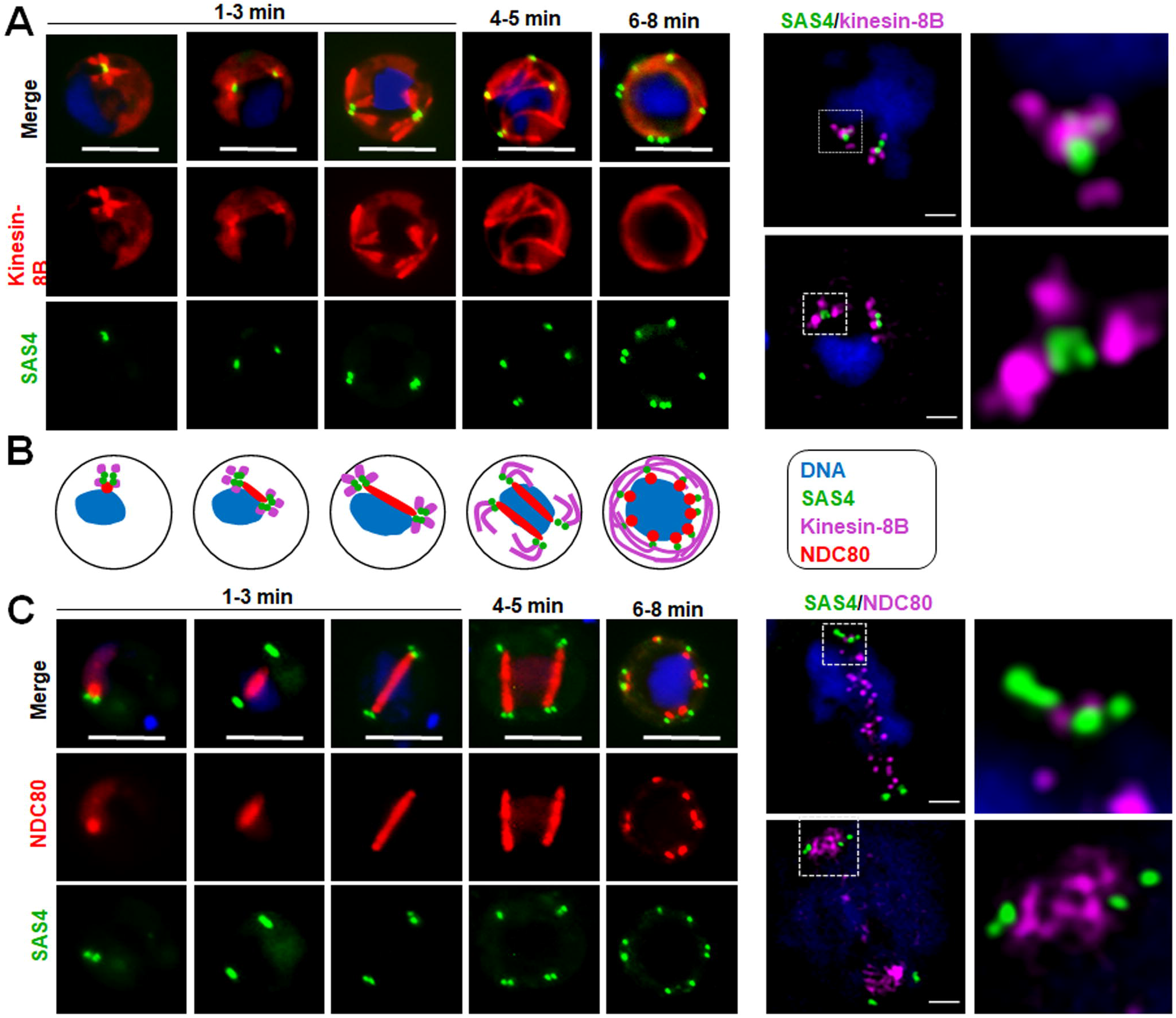
The location of SAS4 in relation to that of the basal body and axoneme (kinesin-8B) and kinetochore (NDC80) markers. **(A)** The location of SAS4-GFP (green) in relation to the basal body and axoneme marker, kinesin-8B-mCherry (red) during male gamete formation. SAS4 shows a cytoplasmic location like kinesin-8B and remains associated with it during basal body biogenesis and axoneme formation throughout male gamete formation. Scale bar = 5 μm. Right hand panel shows super-resolution 3D imaging for SAS4-GFP and kinesin-8B-mCherry localization in gametocytes fixed at 1-2 min post activation. Scale bar =1 μm. (**B)** The location of SAS4-GFP (green) in relation to the kinetochore marker, NDC80-mCherry (red) during male gamete formation. The cytoplasmic location of SAS4 contrasts with the nuclear location of NDC80 during chromosome replication and segregation, indicating that SAS4 is not associated with the mitotic spindle. Scale bar = 5 μm. Right hand panel shows super-resolution 3D imaging for SAS4-GFP and NDC80-mCherry localization in gametocytes fixed at 2-3 min post activation. Scale bar = 1.

### The spatiotemporal locations of SAS4 and the kinetochore protein NDC80 reveal basal body formation and mitotic spindle dynamics are coupled processes

During mitosis in male gametogenesis, genome replication and chromosome segregation are rapid processes. To determine the relationship between mitosis in the nucleus and basal body formation in the cytoplasm, a parasite line expressing both SAS4-GFP, and kinetochore protein NDC80-mCherry was used to image these markers in the same cell. Within 1 mpa, the SAS4-GFP tetrad and the NDC80-mCherry focal point were adjacent but not overlapping close to the nuclear DNA (Fig 2C), with SAS4 in a cytoplasmic location and NDC80 closer to the DNA. Later in gametogenesis the SAS4 tetrad split into two parts with the NDC80 signal extending to form a bridge between them, which is presumably the mitotic spindle decorated with kinetochores (Fig 2C). As the two SAS4 tetrads moved apart the NDC80-positive bridge extended across one side of the nucleus and then separated into two halves (Fig 2C). To resolve further the location of SAS4 tetrads and the NDC80 bridge, we used 3D-SIM on fixed gametocytes expressing these two labelled proteins. The 3D-SIM images clearly showed the two SAS4 tetrads at both ends of the NDC80-positive bridge that then divides into two halves (Fig 2C, right hand panel). The two halves of the NDC80-positive bridge further extend to form two bridges, along with concurrent separation of the SAS4 tetrads into doublets (Fig 2C). This process of NDC80-positive bridge formation and separation continues for a third cycle, resulting in eight NDC80 and SAS4 foci (Fig 2B). During the whole process of NDC80-labelled bridge formation and separation, SAS4 was located adjacent to but never overlapped with NDC80 (Fig 2C). A schematic diagram for this process is provided in Figure 2B.

### Electron microscopy analysis suggests that SAS4 is part of an outer centriolar BB MTOC in male gametocytes

From the literature it is unclear whether the acentriolar MTOC and BB centriolar MTOC are linked and doing similar organisation or there are two independent MTOCs, one that is organising the spindle dynamics in the nucleus and other organising the axoneme biogenesis. Therefore we examined various transmission electron micrographs (TEMs) of male gametocytes and compared them with some of the images of *P yoelii* male gametocytes described earlier by Robert Sinden (Sinden et al., 1976). Our analysis supports the relative location of basal body, axoneme, nucleus and kinetochore, as described by Sinden. In these micrographs, two adjacent electron dense masses are observed on either side of the nuclear membrane (Figs 3A, S2). The outer mass is a typical basal body with nine single α-tubules, whereas the inner part is a nuclear pole. Both structures serve as MTOCs: the basal body for axoneme microtubules (MTs) and the nuclear pole for spindle MTs to which kinetochores are attached (Figs 3A, S2). During mitosis in the asexual blood stages there is no basal body but the MTOC for mitotic spindle MTs is present and located within the nuclear envelope. This observation is consistent with the location of two separate and distinct MTOC. The first one is the nuclear pole (NP) in gametocytes that serves as an inner acentriolar MTOC for spindle MTs. and the second is where SAS4, SAS6 and kinesin-8B are located in the basal body and part of outer centriolar MTOC (Fig 3C). These two independent MTOC have to be coordinated for the successful generation of flagellate male gamete.

**Figure 3.**
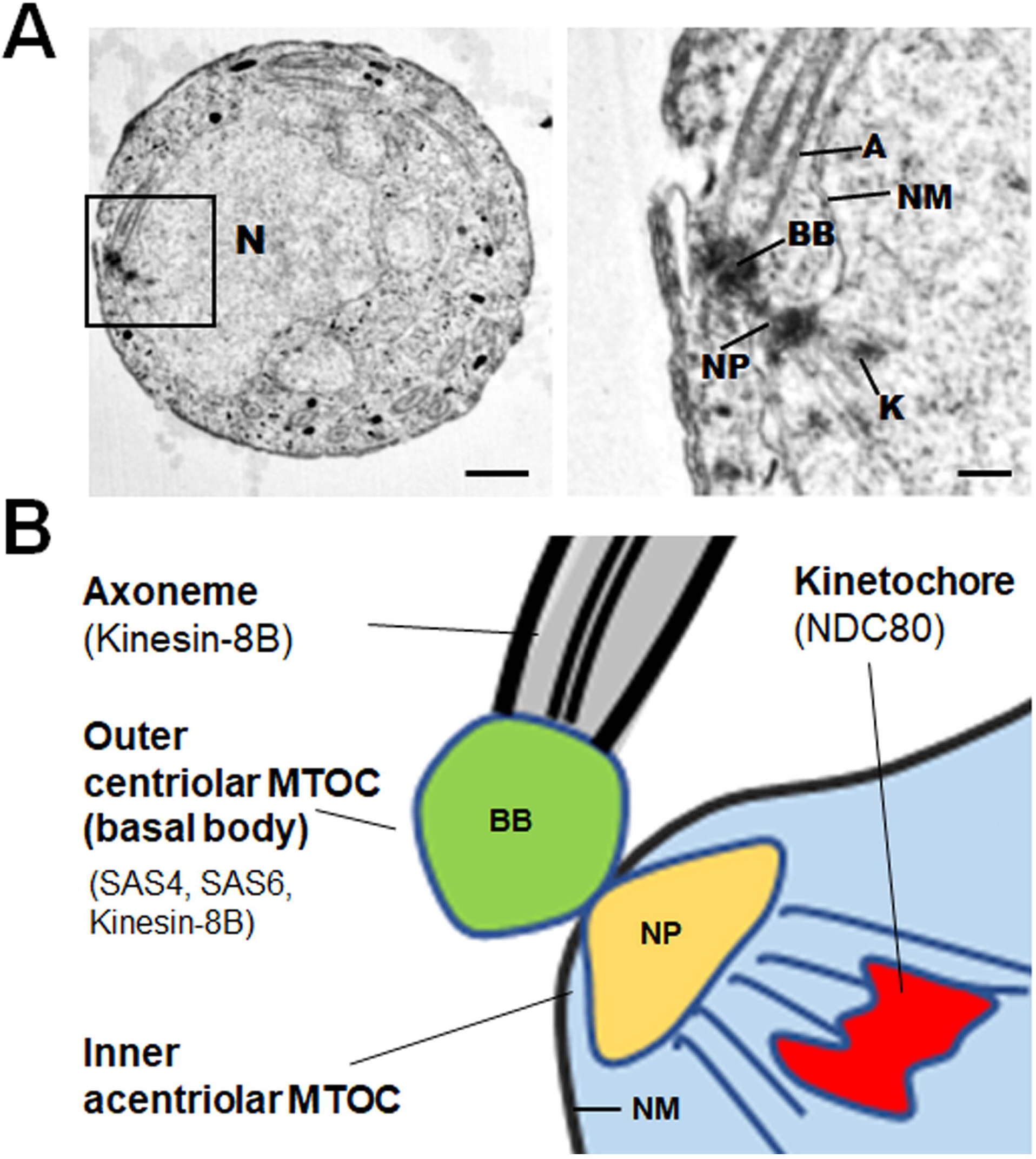
SAS4 is a part of the outer centriolar MTOC (basal body). **(A)** Electron microscopy on 4-5 min post-activation gametocyte reveals the relative locations of basal body, nuclear pole, and kinetochore. Section through the microgametocyte showing a large central nucleus (N) with basal body (BB) in the peripheral cytoplasm. Scale bar = 1 μm. Enlargement of the enclosed area showing the details of each basal body (s) with axoneme in cytoplasmic compartment separated by a nuclear membrane (NM), an intranuclear spindle with attached kinetochores (K) radiating from the nuclear poles (NP). Scale bar = 100nm. **(B)** A schematic diagram showing outer centriolar MTOC (basal body) and inner acentriolar MTOC (NP) serving as microtubule organising centres for axoneme and intranuclear spindle respectively.

### *Plasmodium* SAS4 is dispensable for parasite proliferation and transmission

Based on the expression and location of SAS4 during male gametogenesis and the essential role of basal body protein SAS6 in male gametogenesis (Marques et al., 2015) we examined the importance of SAS4 in male gamete formation. We deleted the gene in a *P. berghei* line constitutively expressing GFP (WT-GFP, Fig S1C) (Janse et al., 2006). Diagnostic PCR showed successful integration of the targeting construct at the *sas4* locus (Fig S1D) and quantitative real time PCR (qRT-PCR) showed the lack of *sas4* expression in gametocytes, confirming the deletion of the *sas4* gene (Fig 4A). Successful creation of the *ΔSAS4* parasite indicated that the gene is not essential in asexual blood stages, consistent with the absence of the protein’s expression at this stage of the life cycle in wild type parasites. Further phenotypic analysis of the *ΔSAS4* parasite was carried in comparison with the parental parasite (WT-GFP).

**Fig 4.**
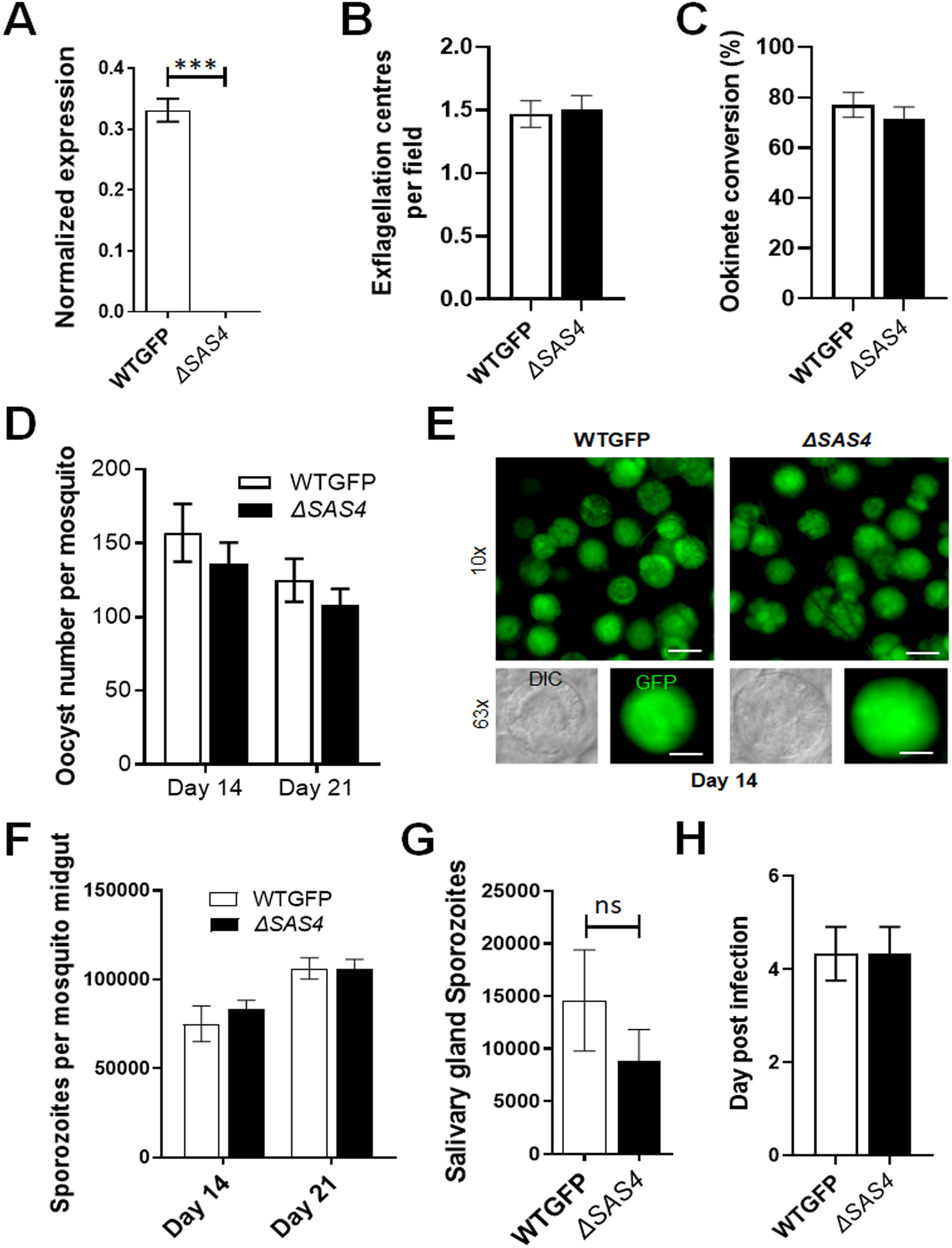
SAS4 is dispensable for parasite proliferation and transmission. **(A)** qRT-PCR analysis of SAS4 transcript in the *ΔSAS4* and WT-GFP parasites to show the complete depletion of *sas4*. **(B)** Male gametogenesis (exflagellation) of *ΔSAS4* parasites compared with WT-GFP parasites measured as the number of exflagellation centres per field. Mean ± SEM. n=3 independent experiments. **(C)** Ookinete conversion as a percentage for *ΔSAS4* and WT-GFP parasites. Ookinetes were identified using 13.1 antibody as a surface marker (P28) and defined as those cells that differentiated successfully into elongated ‘banana shaped’ ookinetes. Mean ± SEM. n=3 independent experiments. **(D)** Total number of GFP-positive oocysts per infected mosquito in *ΔSAS4* compared to WT-GFP parasites at 14 and 21-day post infection. Mean ± SEM. n= 3 independent experiments. **(E)** Mid guts at 10x and 63x magnification showing oocysts of *ΔSAS4* and WT-GFP lines at 14 dpi. Scale bar=50 μM in 10x and 20 μM in 63x. **(F)** Total number of sporozoites in oocysts of *ΔSAS4* and WT-GFP parasites at 14 and 21 dpi. Mean ± SEM. n= 3 independent experiments. **(G)** Total number of sporozoites in salivary glands of *ΔSAS4* and WT-GFP parasites. Bar diagram shows mean ± SEM. n= 3 independent experiments. ns=non-significant **(H)** Bite back experiments showing successful transmission of WT-GFP and *ΔSAS4* parasites from mosquito to mice. Mean ± SEM. n= 3 independent experiments.

First, we examined male gametogenesis, and surprisingly, we observed no significant difference in flagellate gamete formation known as exflagellation in the Δsas4 parasite in comparison with the WT-GFP parasite (Fig 4B). Zygote formation and its differentiation to ookinete development were also unaffected (Fig 4C). To assess the effect of *sas4* gene deletion on oocyst development, the number of GFP-positive oocysts on the mosquito gut wall was counted in mosquitoes fed with either *ΔSAS4-* or WT-GFP parasites, there was no significant difference in the number or size of *ΔSAS4* oocysts compared to WT-GFP controls at 14- and 21-days post-infection (Fig 4D, E). The number of sporozoites produced by *ΔSAS4* and WT-GFP parasites was comparable (Fig 4F), and although there was a slight reduction in numbers of *ΔSAS4* salivary gland sporozoites, the difference from WT-GFP numbers was not significant (Fig 4G). The infectivity of the *ΔSAS4* sporozoites to naïve mice was similar to that of WT-GFP parasites (Fig 4H).

## Discussion

BBs are centriolar organelles that nucleate flagella and cilia, and are important MTOC components, with different and distinct ways of organisation that have arisen during centriole evolution in eukaryotes (Carvalho-Santos et al., 2011; Nabais et al., 2020) SAS6 and SAS4 are core protein components of this organelle (Carvalho-Santos et al., 2010; Hodges et al., 2010). *Plasmodium*, the evolutionarily divergent unicellular eukaryote and causative agent of malaria, shows a rapid and atypical process leading to formation of flagellated male gametes during this stage in its life cycle, within the mosquito gut and a crucial transmission stage. Our recent and earlier ultrastructure studies had identified an amorphous basal body in the cytoplasmic compartment of the male gametocyte, but the properties and function during unusual flagellum formation of SAS4, a conserved CBB molecule, were unknown. During the *Plasmodium* life cycle a centriole is only present during male gametogenesis, whereas during mitosis in other proliferative stages only the amorphous acentriolar MTOC is present (Sinden, 1991). Here we have investigated by live cell imaging in real time the profile of SAS4/CPAP to understand whether it is involved in axoneme biogenesis during male gamete formation and how it’s replication is coordinated during mitosis.

Male gametocyte activation results in rapid genome replication from 1N to 8N in 8 min, with three rounds of mitosis without nuclear division (Sinden, 1991). Our imaging suggests a very rapid de novo formation of SAS4 during male gametogenesis, with a dynamic profile in the cytoplasm. The number of discrete SAS4-GFP foci duplicates as genome replication and rounds of mitosis occur. In most cells, SAS4 appears to coalesce into a close doublet at the beginning of the first mitosis and then this structure replicates in coordination with replication of the genome. We show that eight BB-like structures, each with a close SAS4 doublet are formed de novo and present at the end of genome replication and mitosis. These SAS4 foci appear to be in the cytoplasm of the cell and associated with the outside of the nucleus, suggesting that SAS4 is part of the basal body outer MTOC of. To confirm its location, we generated parasite lines expressing both SAS4-GFP and either the cytoplasmic BB and axoneme marker, Kinesin-8B-mCherry (Zeeshan et al., 2019a), or NDC80-mCherry, a kinetochore marker of the mitotic spindle in the nucleus (Zeeshan et al., 2020b). Real time imaging clearly delineated the spatial organisation of SAS4 with respect to these complementary cytoplasmic and nuclear markers. It was clear that SAS4 is part of the basal body MTOC structure with a similar spatial profile to that of Kinesin-8B. However, SAS4 and Kinesin-8B do not co-localise and the images suggest that SAS4 may be located at the centre of the BB that nucleates axoneme assembly during the male cell differentiation. While Kinesin-8B is part of the axoneme assembly, in contrast SAS4 is limited to the basal body. We show that SAS4 duplication is synchronized with the accumulation of NDC80 and spindle formation during successive rounds of genome replication during mitosis. However, as we have shown previously the NDC80 foci are within the nucleus, our analysis suggests that SAS4 is likely a component of the BB/MTOC in the outer cytoplasmic compartment. We suggest that the MTOC/spindle assembly marked with NDC80 inside the nucleus is coupled together with the cytoplasmic BB/MTOC as cytoskeletal structures. Although *Plasmodium* undergoes closed mitosis, it is possible that these two components of the cell can coordinate mitosis and axoneme assembly to ensure that there is one flagellum for each genome/nucleus at exflagellation. These two components may be inter-connected as part of a bipartite MTOC. A similar structure has been implicated in replication of another apicomplexan parasite, *Toxoplasma gondii* (Suvorova et al., 2015) using asexual cells. Whether biflagellated male gamete of Toxoplasma and Eimeria has similar structure is not known. In Toxoplasma or Eimeria where there is a centriole in the cytoplasm associateed with the nuclear spindle during asexual division, this becomes the basal body at the end of nuclear division to initiate the flagella formation. (Ferguson et al., 1977; Ferguson et al., 1974). In trypanosomes the BB is coupled with kinetoplast DNA during cell division (Vaughan and Gull, 2015) and SAS4 controls the cell cycle transition (Hu et al., 2015), suggesting that the evolution of BBs has depended upon the requirement of the cell to multiply in different niches. Axoneme and flagellum formation only occur during male gametogenesis in *Plasmodium*, and their absence during blood stage schizogony is mirrored by the lack of SAS4 expression at this stage of the life cycle.

We next examined the functional role of SAS4 in *Plasmodium* by examining the phenotype resulting from gene deletion. In contrast to *Plasmodium* SAS6 that was shown to be important for male gamete formation (Marques et al., 2015), we found that SAS4 is not essential for exflagellation as there was no significant change in flagellated gamete formation. At other stages of parasite development, including zygote formation, ookinete and sporozoite development, and parasite transmission and infectivity, no significant differences from the wild type parasite were observed. We conclude that the presence of SAS4 is not essential for parasite development throughout the life cycle. It is possible that other proteins compensate for SAS4 function, or it has a redundant function in *Plasmodium*.

It will be interesting in the future to analyse in depth the 3D structure of *Plasmodium* BBs and their components, for example by using these BB and mitotic spindle markers and correlative light and electron microscopy (CLEM) as described recently for the bryophyte Physocomistrium (Gomes Pereira et al., 2021). In a recent study Raspa and Brochet have used expansion microscopy to study the MTOC structures in Plasmodium (Rashpa and Brochet, 2021) The protein interactome obtained using these lines and others will provide tools to identify BB components and understand their evolution in this divergent organism, which assembles the complete BB complement de novo in eight minutes. This is extremely fast, for example when compared to the rapid assembly that takes one hour in Naegleria (Fritz-Laylin et al., 2016). These approaches may also identify the conserved and divergent molecules that enable the extremely fast flagellum assembly, which is one of the fastest known and where accuracy may be compromised because of the need for speed. (Fritz-Laylin et al., 2016; Sinden et al., 2010).

Overall, this study shows that SAS4 is part of an outer cytoplasmic BB MTOC where there is a need for coordination between flagellum assembly in the cytoplasm and genome replication in the nucleus so that there is one flagellum for each haploid nucleus formed following karyokinesis. Formation of the BB occurs de novo and the entire process is very rapid. However, the deletion of the SAS4 gene does not affect male gametogenesis and the gene is not essential for parasite transmission or at other stages of the life cycle.

## Materials and Methods

### Ethics statement

The animal work passed an ethical review process and was approved by the United Kingdom Home Office. Work was carried out under UK Home Office Project Licenses (30/3248 and PDD2D5182) in accordance with the United Kingdom ‘Animals (Scientific Procedures) Act 1986’. Six-to eight-week-old female CD1 outbred mice from Charles River laboratories were used for all experiments.

### Generation of transgenic parasites

The C-terminus of SAS4 was tagged with GFP by single crossover homologous recombination in the parasite. To generate the SAS4-GFP line, a region of the *sas4* gene downstream of the ATG start codon was amplified using primers T2011 and T2012, ligated to p277 vector, and transfected as described previously (Saini et al., 2017). A schematic representation of the endogenous *sas4* locus (PBANKA_1322200), the constructs and the recombined *sas4* locus are shown in Fig S1A. The oligonucleotides used to generate the mutant parasite lines are described in Table S1. *P. berghei* ANKA line 2.34 (for GFP-tagging) or ANKA line 507cl1 expressing GFP (for gene deletion) were transfected by electroporation (Janse et al., 2006)

The gene-deletion targeting vector for *sas4 was* constructed using the pBS-DHFR plasmid, which contains polylinker sites flanking a *T. gondii dhfr/ts* expression cassette conferring resistance to pyrimethamine, as described previously (Zeeshan et al., 2019a). PCR primers N1391 and N1392 were used to generate a 803 bp fragment of *sas4* 5′ upstream sequence from genomic DNA, which was inserted into *Apa*I and *Hin*dIII restriction sites upstream of the *dhfr/ts* cassette of pBS-DHFR. A 721 bp fragment generated with primers N1393 and N1394 from the 3′ flanking region of *sas4* was then inserted downstream of the *dhfr/ts* cassette using *Eco*RI and *Xba*I restriction sites. The linear targeting sequence was released using *Apa*I/*Xba*I. A schematic representation of the endogenous *sas4* locus the constructs and the recombined *sas4* locus can be found in Fig S1C.

### Parasite genotype analyses

For the parasites expressing a C-terminal GFP-tagged SAS4 protein, diagnostic PCR was used with primer 1 (IntT201) and primer 2 (ol492) to confirm integration of the GFP targeting construct (Fig S1B). For the gene knockout parasites, diagnostic PCR was used with primer 1 (IntN139) and primer 2 (ol248) to confirm integration of the targeting construct, and primer 3 (N139 KO1) and primer 4 (N139 KO2) were used to confirm deletion of the *sas4* gene (Fig S1D).

### Purification of gametocytes

The purification of gametocytes was achieved using a protocol described previously (Beetsma et al., 1998) with some modifications. Briefly, parasites were injected into phenylhydrazine treated mice and enriched by sulfadiazine treatment after 2 days of infection. The blood was collected on day 4 after infection and gametocyte-infected cells were purified on a 48% v/v NycoDenz (in PBS) gradient. (NycoDenz stock solution: 27.6% w/v NycoDenz in 5 mM Tris-HCl, pH 7.20, 3 mM KCl, 0.3 mM EDTA). The gametocytes were harvested from the interface and washed.

### Live cell- and -time lapse imaging

Purified gametocytes were examined for GFP expression and localization at different time points (1-15 min) after activation in ookinete medium containig xanthurenic acid. Images were captured using a 63x oil immersion objective on a Zeiss Axio Imager M2 microscope fitted with an AxioCam ICc1 digital camera (Carl Zeiss, Inc). Time-lapse videos (1 frame every 5 sec for 10 cycles) were taken with a 63x objective lens on the same microscope and analysed with the AxioVision 4.8.2 software as described recently (Zeeshan et al., 2020b).

### Generation of dual tagged parasite lines

The SAS4-GFP parasites were mixed with kinesin-8B-mCherry or NDC80-mCherry parasites in equal numbers and injected into a mouse. Mosquitoes were fed on this mouse 4 to 5 days after infection when gametocyte parasitaemia was high. These mosquitoes were checked for oocyst development and sporozoite formation at day 14 and day 21 after feeding. Infected mosquitoes were then allowed to feed on naïve mice and after 4 - 5 days these mice were examined for blood stage parasitaemia by microscopy with Giemsa-stained blood smears. In this way, some parasites expressed both SAS4-GFP and kinesin-8B-mCherry or SAS4-GFP and NDC80-mCherry in the resultant gametocytes. These gametocytes were purified, and fluorescence microscopy images were collected as described above.

### Super resolution microscopy

A small volume (3 μl) of gametocytes was mixed with Hoechst dye and pipetted onto 2 % agarose pads (5×5 mm squares) at room temperature. After 3 min these agarose pads were placed onto glass bottom dishes with the cells facing towards glass surface (MatTek, P35G-1.5-20-C). Cells were scanned with an inverted microscope using Zeiss C-Apochromat 63×/1.2 W Korr M27 water immersion objective on a Zeiss Elyra PS.1 microscope, using the structured illumination microscopy (SIM) technique. The correction collar of the objective was set to 0.17 for optimum contrast. The following settings were used in SIM mode: lasers, 405 nm: 20%, 488 nm: 50%; exposure times 100 ms (Hoechst) and 25 ms (GFP); three grid rotations, five phases. The band pass filters BP 420-480 + LP 750 and BP 495-550 + LP 750 were used for the blue and green channels, respectively. Multiple focal planes (Z stacks) were recorded with 0.2 μm step size; later post-processing, a Z correction was done digitally on the 3D rendered images to reduce the effect of spherical aberration (reducing the elongated view in Z; a process previously tested with fluorescent beads). Images were processed and all focal planes were digitally merged into a single plane (Maximum intensity projection). The images recorded in multiple focal planes (Z-stack) were 3D rendered into virtual models and exported as images. Processing and export of images were done by Zeiss Zen 2012 Black edition, Service Pack 5 and Zeiss Zen 2.1 Blue edition (Zeeshan et al., 2020b).

### Parasite phenotype analyses

Blood containing approximately 50,000 parasites of the *ΔSAS44* line was injected intraperitoneally (i.p) into mice to initiate infections. Asexual stages and gametocyte production were monitored by microscopy on Giemsa-stained thin smears. Four to five days post infection, exflagellation and ookinete conversion were examined as described previously(Zeeshan et al., 2019b) with a Zeiss AxioImager M2 microscope (Carl Zeiss, Inc) fitted with an AxioCam ICc1 digital camera. To analyse mosquito transmission, 30–50 *Anopheles stephensi* SD 500 mosquitoes were allowed to feed for 20 min on anaesthetized, infected mice with an asexual parasitemia of 15% and a comparable number of gametocytes as determined on Giemsa-stained blood films. To assess mid-gut infection, approximately 15 guts were dissected from mosquitoes on day 14 post feeding, and oocysts were counted on an AxioCam ICc1 digital camera fitted to a Zeiss AxioImager M2 microscope using a 63x oil immersion objective. On day 21 post-feeding, another 20 mosquitoes were dissected, and their guts crushed in a loosely fitting homogenizer to release sporozoites, which were then quantified using a haemocytometer. Mosquito bite back experiments were performed 21 days post-feeding using naive mice, and blood smears were examined after 3-4 days.

### Electron microscopy

Gametocytes activated for 4-5 min were fixed in 4% glutaraldehyde in 0.1 M phosphate buffer and processed for electron microscopy as previously described (Zeeshan et al., 2019b). Briefly, samples were post fixed in osmium tetroxide, treated *en bloc* with uranyl acetate, dehydrated and embedded in Spurr’s epoxy resin. Thin sections were stained with uranyl acetate and lead citrate prior to examination in a JEOL1200EX electron microscope (Jeol UK Ltd).

### Quantitative Real Time PCR (qRT-PCR) analyses

RNA was isolated from gametocytes using an RNA purification kit (Stratagene). cDNA was synthesised using an RNA-to-cDNA kit (Applied Biosystems). Gene expression was quantified from 80 ng of total RNA using a SYBR green fast master mix kit (Applied Biosystems). All the primers were designed using the primer3 software (Primer-blast, NCBI). Analysis was conducted using an Applied Biosystems 7500 fast machine with the following cycling conditions: 95°C for 20 s followed by 40 cycles of 95°C for 3 s; 60°C for 30 s. Three technical replicates and three biological replicates were performed for each assayed gene. The *hsp70* (PBANKA_081890) and *arginyl-t RNA synthetase* (PBANKA_143420) genes were used as endogenous control reference genes. The primers used for qPCR can be found in **Table S1.**

### Statistical analysis

All statistical analyses were performed using GraphPad Prism 7 (GraphPad Software). For qRT-PCR, an unpaired t-test was used to examine significant differences between wild-type and mutant strains.

## Acknowledgement

We thank the Oxford Brookes Centre for Bioimaging for assistance.

## Supplementary materials

### Supplementary Figures

**Fig S1. Generation and genotypic analysis of SAS4-GFP and *ΔSAS4* parasites (A)** Schematic representation of the endogenous *Pbsas4* locus, the GFP-tagging construct and the recombined *sas4* locus following single homologous recombination. Arrows 1 and 2 indicate the position of PCR primers used to confirm successful integration of the construct. **(B)** Diagnostic PCR of *SAS4-GFP* (tag) and WT parasites using primers IntT201 (Arrow 1) and ol492 (Arrow 2). Integration of the sas4 tagging construct gives a band of 1159 bp. **(C)** Schematic representation of *Pbsas4* locus, knockout construct and the recombined sas4 locus following double homologous recombination. Arrows 1 (intN139) and 2 (ol248) indicate the primers position used to confirm 5’ integration and arrows 3 (N139KO1) and 4 (N139KO1) indicate the primers used to check the deletion of gene **(D)** Integration PCR of the *sas4* locus in WTGFP (WT) and knockout (Mut) parasites showing expected band size of integration in knockout parasites and knockout (KO) PCR showing deletion from mut parasites.

**Fig S2. Electron micrographs of gametocyte reveals the relative locations of basal body, nuclear pole, and kinetochore**

Enlarged electron micrographs of gametocytes activated for 4-5 min showing the details of basal body (bb) with axoneme (A) in cytoplasmic compartment separated by a nuclear membrane (NM), an intranuclear spindle with attached kinetochores (K) radiating from the nuclear poles (NP). Scale bar = 100nm.

### Supplementary Tables

**Table S1.** Oligonucleotides used in this study.

### Supplementary Videos

**Video S1.** Video for SAS4-GFP localization in male gametocytes at 1-2 min post activation

**Video S1.** Video for SAS4-GFP localization in male gametocytes at 3-4 min post activation

