## Supplementary figures and images for "*Plasmodium* SAS4/CPAP is a flagellum basal body component during male gametogenesis, but is not essential for parasite transmission"

### Fig S1

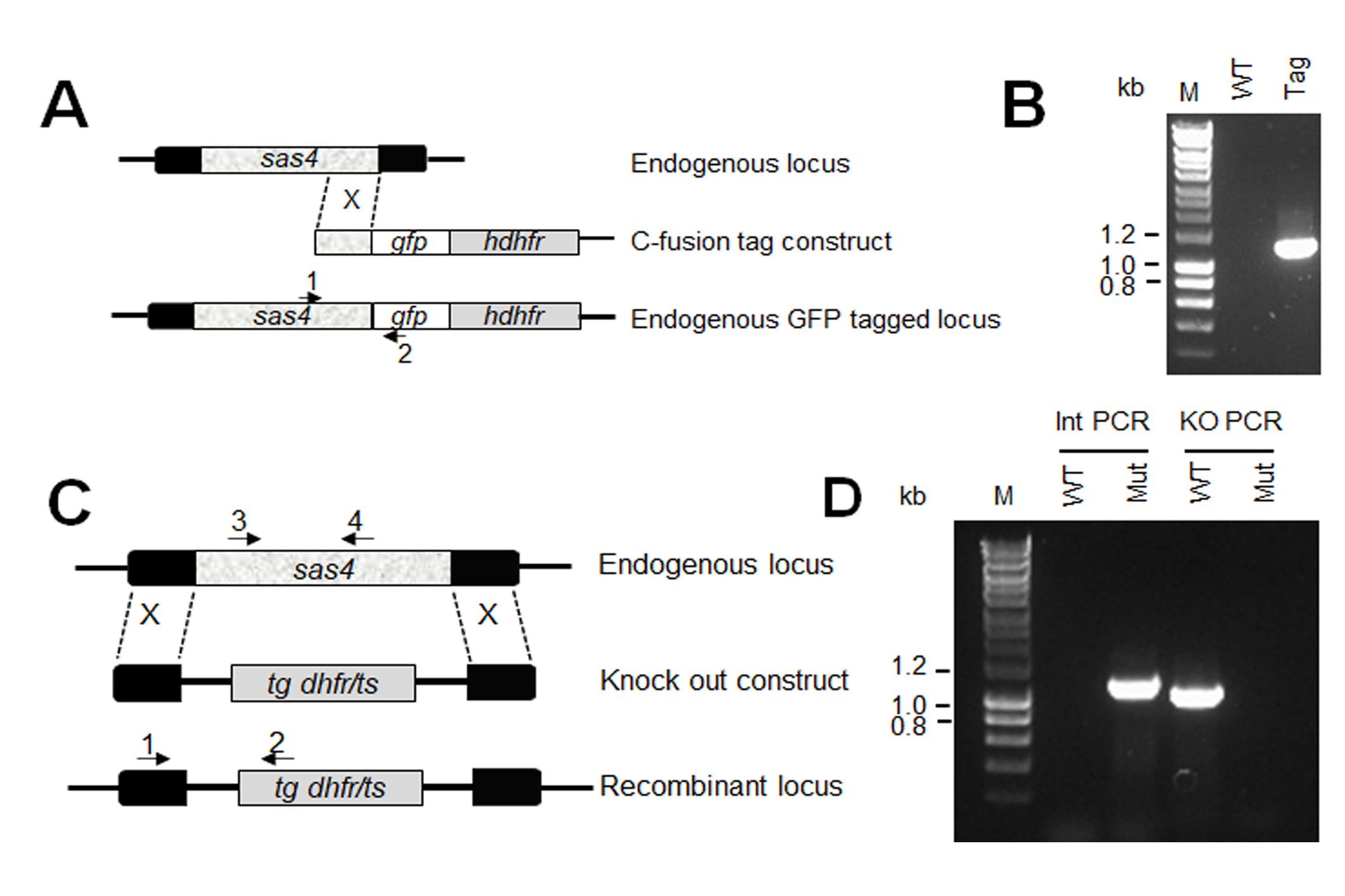

### Fig S2

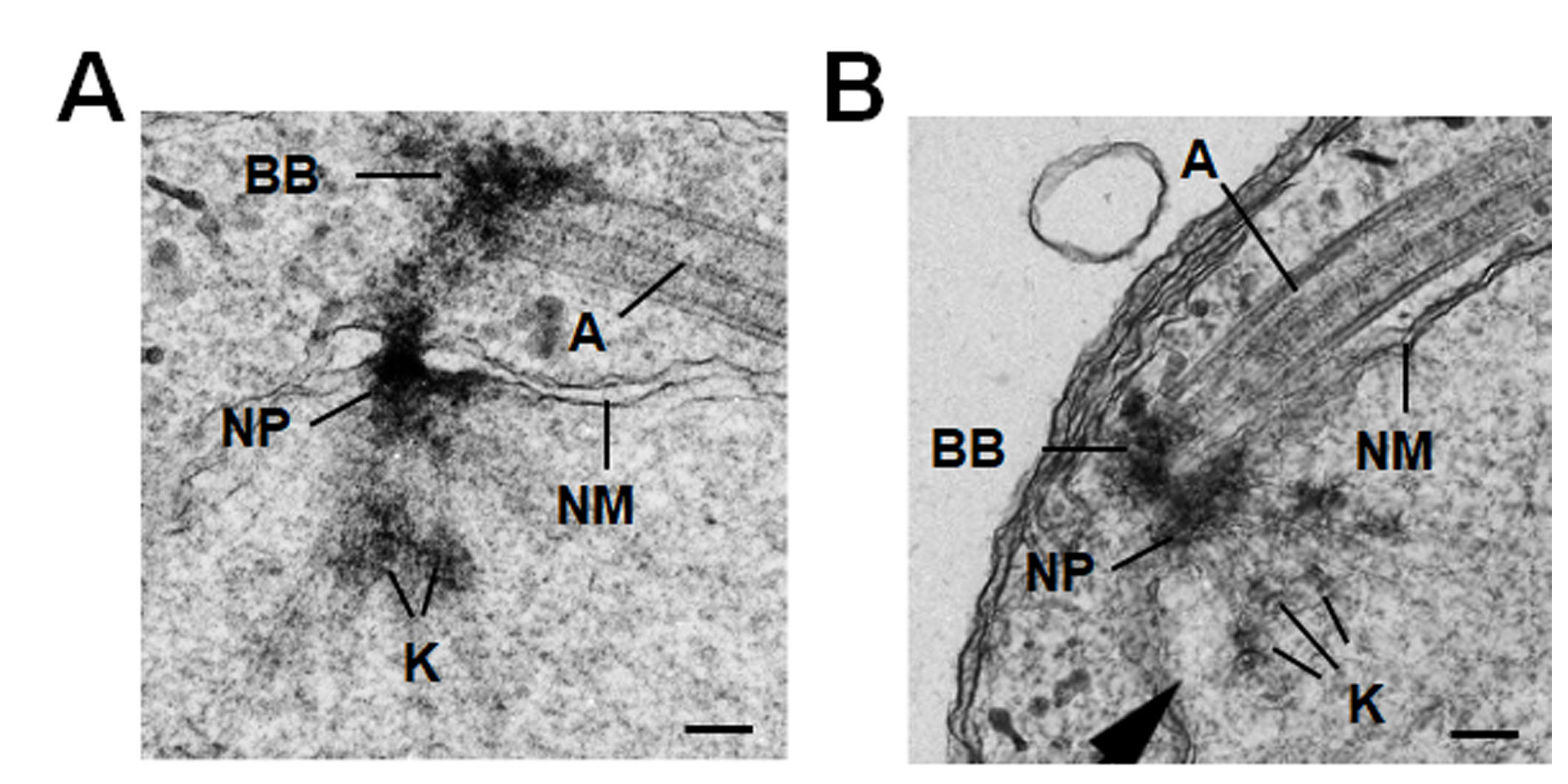
